# Unifying the two different classes of plant non-specific lipid-transfer proteins allergens classified in the WHO/IUIS allergen database through a motif with conserved sequence, structural and electrostatic features

**DOI:** 10.1101/087411

**Authors:** Sandeep Chakraborty

## Abstract

The ubiquitously occuring non-specific lipid-transfer proteins (nsLTPs) in plants are implicated in key processes like biotic and abiotic stress, seed development and lipid transport. Additionally, they constitute a panallergen multigene family present in both food and pollen. Presently there are 49 nsLTP entries in the WHO/IUIS allergen database (http://allergen.org/). Analysis of full-length allergens identified only two major classes (nsLTP1,n=32 and nsLTP2,n=2), although nsLTPs are classified into many other groups. nsLTP1 and nsLTP2 are differentiated by their sequences, molecular weights, pattern of the conserved disulphide bonds and volume of the hydrophobic cavity. The conserved R44 is present in all full length nsLTP1 allergens (only Par j 2 from *Parietaria judaica* has K44), while D43 is present in all but Par j 1/2 from *P. judaica* (residue numbering based on PDBid:2ALGA). Although, the importance of these residues is well-established in nsLTP1, the corresponding residues in nsLTP2 remain unknown. A structural motif comprising of two cysteines with a disulphide bond (C3-C50), R44 and D43 identified a congruent motif (C3/C35/R47/D42) in a nsLTP2 protein from rice (PDBid:1L6HA), using the CLASP methodology. This also provides a quantitative method to assess the cross-reactivity potential of different proteins through congruence of an epitope and its neighbouring residues. Future work will involve obtaining the PDB structure of an nsLTP2 allergen and Par j 1/2 nsLTP1 sequences with a missing D43, determine whether nsLTP from other groups beside nsLTP1/2 are allergens, and determine nsLTP allergens from other plants commonly responsible for causing allergic reactions (chickpea, walnut, etc.) based on a genome wide identification of genes with conserved allergen features and their *in vitro* characterization.

## Introduction

IgE-mediated food and pollen allergy are manifested with severe clinical symptoms, and is a rapidly growing worldwide health concern [1]. ‘The WHO/IUIS Allergen Nomenclature Sub-committee is responsible for maintaining and developing a unique, unambiguous and systematic nomenclature for allergenic proteins’ and ‘maintains an allergen database that contains approved and officially recognized allergens’ (http://allergen.org/) [2]. Allergens are typically restricted to a few classes of proteins, possessing similar biochemical functions [3]. Plant non-specific lipid-transfer proteins (nsLTP) are an important panallergen family [4] of both food and pollen allergens (with 49 entries in the database currently) [5]. nsLTPs are involved in key processes, such as the stabilization of membranes [6], resistance to biotic [7–9] and abiotic stress [10], long distance signaling [11], sexual reproduction [12], seed development [13] and germination [14]. nsLTPs are extremely resistant to heat and proteolytic digestion [15], sensitized by inhalation or ingestion, and implicated with systemic and severe allergic symptoms (rhinitis, conjunctivitis, dermatitis, asthma and anaphylaxis) [16].

nsLTPs belong to the PR-14 pathogenesis related protein, and share several characteristics (basic, <10kDA, conserved four disulphide bonds of a eight-cysteine motif (8C) [18]) [17]. Several classifications of nsLTPs have been proposed based on the spacing of the 8C spacing [19–21]. Interestingly, all documented allergen nsLTPs are limited to two classes - nsLTP1 and nsLTP2 [22]. nsLTP1 and nsLTP2 are di erentiated by their sequences, molecular weights, pattern of the conserved disulphide bonds and volume of the hydrophobic cavity [19]. There is no solved structure for a nsLTP2 allergen, while several structures exist for nsLTP1 allergens. The number of full length nsLTP1s in the allergen database far exceeds the number of nsLTP2s [22]. Also, the epitope R44 for nsLTP1 has been determined [23, 24]. While R44 is absolutely conserved in all full-length nsLTP1 allergens, the corresponding residues have not been determined in nsLTP2 allergens.

Epitope prediction accelerates the determination of peptides that bind to IgE [23]. Viral epitopes can be identified by perturbations in the envelope protein that mediates viral fusion with a host cell [25,26]. Similar perturbations in nsLTP on lipid binding renders them susceptible to proteolysis [27]. Thus, perturbation analysis might not be a good strategy for epitope prediction in nsLTPs. The spatial and electrostatic congruence of active site residues in proteins with the same functionality, even those convergently evolved like serine proteases [28], has been demonstrated on previous occasions [29–33]. A similar strategy can be adopted for allergen epitopes by using a motif from a known allergen (such as R44 in nsLTP1, PDBid:2ALGA) to query other structures from nsLTP2 with unknown epitopes.

Here, nsLTP allergens (n=49) from the allergen database (http://allergen.org/) are analyzed. Fragments (<40aa, n=6) are removed, and those with known Uniprot ids (n=34) are grouped using the YeATS suite [34] to identify two main classes (nsLTP1 and nsLTP2), corroborating previous results [22]. Moreover, the fragment allergen Ole e 7 is not homologous to any other nsLTP, and is probably mis-annotated. A motif obtained from nsLTP1 (PDBid:2ALGA, the prototypical peach Pru p 3 [35]) using conserved residues is used to query a nsLTP2 structure (PDBid:1L6HA from rice [36], homologous to the two known nsLTP2 allergens) using CLASP [29]. The presence of a congruent motif in nsLTP2 indicates that this might be the corresponding epitope in nsLTP2, although this would require experimental validation.

## Results and discussion

### Classifying nsLTP allergens in http://allergen.org/

Two keyword searches - ‘lipid’ and ‘nsLTP’- were used to query the http://allergen.org/, giving 44 and 22 results, respectively. These were merged to obtain 49 nsLTP allergens (Table 1). 40 sequences have Uniprot ids assigned, and 6 sequences are fragments (<40 aa). The fragmented allergen Ole e 7 (Uniprot id:P81430, length=21) is not homologous to any other nsLTP, and is probably mis-annotated (SI Fig. 1). 34 full length protein sequences were analyzed using the ‘YeATS-GROUP’ algorithm (see Methods). This grouped 32 sequences as nsLTP1, and 2 sequences as nsLTP2 with BLAST bitscore (BBS) =60. nsLTP1 and nsLTP2 are di erentiated by their sequences (<30%identity), molecular weights (9kDa and 7kDa, respectively), pattern of the conserved disulphide bonds and volume of the hydrophobic cavity (nsLTP1 has a larger cavity) [19].

**Table 1:** Cataloguing 49 nsLTPs in the allergen.org database: Allergens that do not have a keyword “lipid” are postfixed with a * - these allergens were extracted using the keyword “nsLTP”. The PDB structure of the allergen Cor a 8 (PDBid: 4XUW) is not annotated in the website. The fragment Ole e 7 is not homologous to any other nsLTP, and is probably mis-annotated.

| Allergen name | Uniprot ids | Length | PDB ids |
| --- | --- | --- | --- |
| Pru p 3 | P81402 Q9LED1 | 91 | 2ALG,2B5S |
| Zea m 14 | P19656-1 P19656-2 | 120 | 1AFH,1MZL,1FK0 |
| Cor a 8 | Q9ATH2 | 115 | 4XUW* |
| *Pis s 3 | C0HJR7 | 120 | 2N81 |
| Len c 3 | A0AT29 | 118 | 2MAL |
| Amb a 6 | O04004 | 118 |  |
| Api g 2 | E6Y8S8 | 118 |  |
| Api g 6 | P86809 | 67 |  |
| Ara h 9 | B6CEX8 B6CG41 | 116 |  |
| Can s 3 | W0U0V5 | 91 |  |
| Fra a 3 | Q8VX12 Q4PLT9 Q4PLU0 Q4PLT6 | 117 |  |
| Hel a 3 | Q7X9Q5 | 116 |  |
| Hev b 12 | Q8RYA8 | 116 |  |
| Jug r 3 | C5H617 | 119 |  |
| Mal d 3 | Q5J026 Q5J011 Q5J009 Q5IZZ6 Q5IZZ5 | 115 |  |
| Mor n 3 | P85894 | 91 |  |
| Par j 1 | P43217 O04404 Q1JTN5 Q40905 | 139 |  |
| Par j 2 | P55958 O04403 | 133 |  |
| Pha v 3 | D3W146 D3W147 | 115 |  |
| Pru ar 3 | P81651 | 91 |  |
| Pru av 3 | Q9M5X8 | 117 |  |
| Pru d 3 | P82534 | 91 |  |
| Pru du 3 | C0L0I5 | 123 |  |
| Pun g 1 | A0A059STC4 A0A059SSZ0 A0A059ST23 | 120 |  |
| Pyr c 3 | Q9M5X6 | 115 |  |
| Rub i 3 | Q0Z8V0 | 117 |  |
| Sin a 3 | E6Y2L9 | 92 |  |
| Sola l 3 | P93224 | 114 |  |
| Sola l 6 | K4BBD9 | 94 |  |
| Tri a 14 | D2T2K2 | 92 |  |
| Vit v 1 | Q850K5 | 119 |  |
| Cit s 3 | Q6EV47 P84161 | 91 |  |
| *Pla or 3 | A9YUH6 | 118 |  |
| *Act d 10 | P86137 P85206 | 92 |  |
| Cit r 3 | P84161 | 20 |  |
| Cit l 3 | P84160 | 20 |  |
| Mus a 3 | P86333 | 39 |  |
| Art v 3 | P0C088 C4MGG9 C4MGH0 C4MGH1 | 37 |  |
| *Act c 10 | P85204 | 15 |  |
| X Ole e 7 | P81430 | 21 |  |
| Pla a 3 |  |  |  |
| Ara h 16 |  |  |  |
| Ara h 17 |  |  |  |
| Aspa o 1 |  |  |  |
| Bra o 3 |  |  |  |
| Cas s 8 |  |  |  |
| Lac s 1 |  |  |  |
| Par o 1 |  |  |  |
| *Sola l 7 |  |  |  |

**Figure 1:**
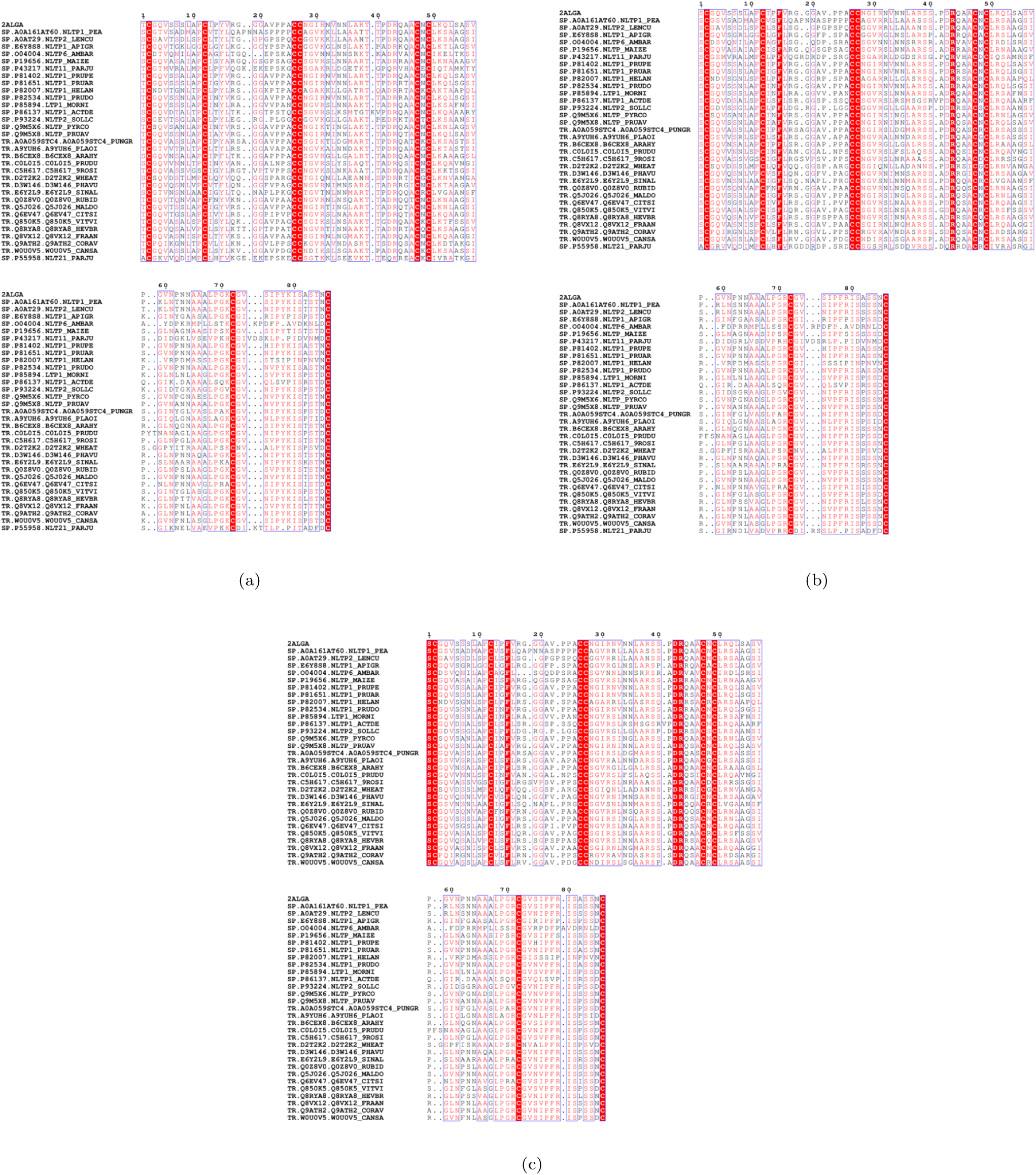
Multiple sequence alignment of nsLTP1 sequences in the allergen.org database: Residue numbering is based on the structure (PDBid:2ALGA) of the prototypical peach Pru p 3. **(a)** Only the 8 cysteine residues forming 4 disulphide bonds are completely conserved. **(b)** Replacing stereochemically equivalent residues identified an aromatic residue (Y16 or F16) and a basic residue (R44 or K44) as being conserved in all sequences. D43 is missing in two allergens from *Parietaria judaica* (sticky-weed, Par j 1 and 2), which is possibly compensated by glutamic acid residues around R44. **(c)** Removing Par j 1 and 2 allergens from sticky-weed, and replacing stereochemically equivalent residues identifies D43 - and another serine or threonine (S2/T2) - as being conserved.

### nsLTP1 - creating a structural motif

The multiple sequence alignment of the nsLTP1 sequences demonstrates the conservation of the cysteine residues, but shows no other absolute conservation among residues (Fig 1a). Replacing for equivalent residues (see Methods) highlights an aromatic residue (Y16 or F16) and a basic residue (R44, K44 only in Par j 2 [37], from *Parietaria judaica*) being common among all sequences (Fig 1b). Residue numbering is based on the structure (PDBid:2ALGA) of the prototypical peach Pru p 3 [35]. Furthermore, Par j 2 and another allergen from the same sticky-weed plant (Par j 1) do not have a conserved D43 residue, contradicting the statement that ‘both D43 and R44 are strictly conserved in LTPs’ [38]. The missing D43 in these two sequences is possibly compensated by extra glutamic acid residues, not found in other sequences, on both sides of R44. Since, the presence of glutamic acid induces disorder in protein structures [39], it would be interesting to see the di erences in the structures of these proteins with other solved nsLTP1s. Removing Par j 1 and 2 sequences, and replacing for equivalent residues highlights another additional conserved residue (Ser2/Thr2) (Fig 1c). Serine and threonine are stereochemically equivalent - for example, N-Linked glycosylation usually occurs at Asn-X-Ser/Thr. While, conversation does not imply that a residue will be an epitope (since it may have structural relevance like the 8C motif), it provides a frame of reference for predicted epitopes.

A motif using D43/R44 and two cysteines from the 8C configuration (Cys3-Cys50) was used to create a four residue motif (nsLTP1Motif:C3/C50/R44/D43). CLASP queried several solved structures of these nsLTP1 allergens using nsLTP1Motif (Table 2). The pairwise distance and electrostatic potential di erence (EPD) in these residues show that although there is overall similarity, there are certain di erences in distances and EPD. For example, even for the same nsLTP from maize, the distance between C52SG and D45OD1 varies from 9.8 Å in PDBid:1MZLA to 13.9 Å in PDBid:1AFHA (both structures are ligand free). While, this di erence might be experimental error, it is hypothesized that in general these di erences might correlate with allergenicity and cross-reactivity. It is not straightforward to identify the residues corresponding to nsLTP1Motif in nsLTP2s due to low sequence homology.

**Table 2:** Potential and spatial congruence of conserved residues in all nsLTP allergens demonstrated through CLASP: The ordering of residues forming disulphide bonds are di erent in nsLTP1 and nsLTP2. Two cysteines (C3-C50) are combined with the conserved R46 and D45 (numbering based on PDBid:1AFHA) to create a motif (nsLTP1Motif). The cognate residues in other nsLTP1 proteins, though mostly similar, show some di erences. nsLTP1Motif also identified a conguent configuration in the nsLTP2 structure from rice. The following atoms were used for the amino acids - C:SG, R:NH1 and D:OD1. DIST = Pairwise distance in Å PD = Pairwise potential di erence. See Methods section for units of potential.

| Type | Method | Name PDBid | nsLTP1Motif(a,b,c,d) |  | ab | ac | ad | bc | bd | cd |
| --- | --- | --- | --- | --- | --- | --- | --- | --- | --- | --- |
| nsLTP1 | *2.3 Å | Peach 2ALGA | C3,C50,R44,D43, | DIST | 2 | 15 | 11.2 | 12.9 | 9.7 | 10 |
|  |  |  |  | PD | 23.9 | 109.4 | 134.5 | 85.5 | 110.6 | 25.1 |
|  | 2.4 Å | Peach 2B5SA | C3,C50,R44,D43, | DIST | 2 | 14.9 | 10.9 | 12.8 | 9.5 | 10 |
|  |  |  |  | PD | 49.2 | 105 | 310.2 | 55.8 | 261 | 205.2 |
|  | NMR | Lentil 2MALA | C4,C51,R45,D44, | DIST | 2 | 18.2 | 13.3 | 16.3 | 11.9 | 11 |
|  |  |  |  | PD | -57.1 | 144 | 134 | 201.1 | 191.1 | -10 |
|  | NMR | Pea 2N81A | C4,C53,R47,D46, | DIST | 2 | 16.6 | 11.4 | 14.9 | 11 | 11.2 |
|  |  |  |  | PD | -19.4 | 49.3 | 132.2 | 68.7 | 151.7 | 83 |
|  | *1.1 Å | Hazelnut 4XUWA | C27,C74,R68,D67, | DIST | 2.1 | 17.3 | 12.7 | 15.9 | 12.1 | 8 |
|  |  |  |  | PD | -20.7 | 220 | 215.7 | 240.7 | 236.4 | -4.3 |
|  | NMR | Maize 1AFHA | C4,C52,R46,D45, | DIST | 2 | 17.3 | 13.8 | 16.3 | 13.9 | 9.2 |
|  |  |  |  | PD | -31.3 | 76.6 | 176.2 | 107.9 | 207.5 | 99.6 |
|  | 1.9 Å | Maize 1MZLA | C4,C52,R46,D45, | DIST | 2 | 16.8 | 11.2 | 14.9 | 9.8 | 9.2 |
|  |  |  |  | PD | 26.6 | 189.1 | 236.9 | 162.5 | 210.3 | 47.7 |
|  | *1.8 Å | Maize 1FK0A | C4,C52,R46,D45, | DIST | 2.1 | 17.4 | 13.2 | 15.6 | 11.5 | 7.9 |
|  |  |  |  | PD | -31 | 134.6 | 169.4 | 165.6 | 200.4 | 34.8 |
| nsLTP2 |  | Rice 1L6HA | C3,C35,R47,D42, | DIST | 2 | 15.8 | 11.7 | 17.6 | 13.3 | 8.7 |
|  |  |  |  | PD | 0 | 71.6 | 336.9 | 71.5 | 336.9 | 265.3 |
|  | NMR | Invert disulphide bond | C35,C3,R47,D42, | DIST | 2 | 17.6 | 13.3 | 15.8 | 11.7 | 8.7 |
|  |  |  |  | PD | -0 | 71.5 | 336.9 | 71.6 | 336.9 | 265.3 |

### nsLTP2 - searching the structure using the nsLTP1Motif

There are no solved structures for the two nsLTP2 allergens (Fig 2a). The structure of an homologous nsLTP2 from rice (PDBid:1L6HA) was used as the representative of nsLTP2, although the protein is not a documented allergen. PDBid:1L6HA was queried using nsLTP1Motif, revealing the presence of a homologous configuration with similar electrostatic and spatial features (C3/C35/R47/D42) (Table 2). The nsLTP1Motif in maize nsLTP1 (PDBid:1FK0A) has the highest congruence with nsLTP2 in rice, and the congruence increases if the cysteine residues having the disulphide bond (C3-C35) are inverted. Note, that R47 in nsLTP2 is not exactly aligned and is shifted by one bit (Fig 2a), which can be fixed by manual insertion of a single gap (Fig 2b).

**Figure 2:**
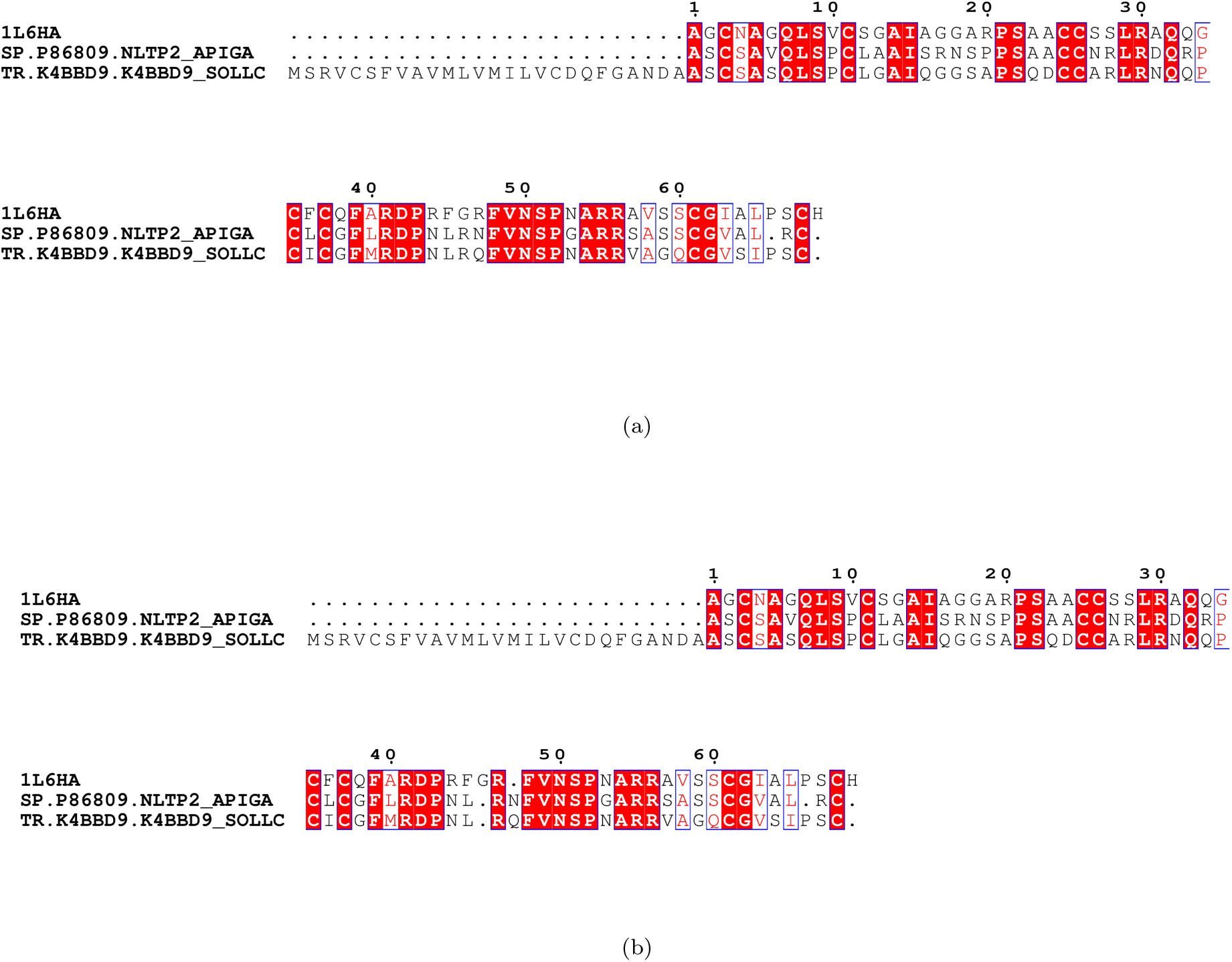
Multiple sequence alignment of nsLTP2 sequences in the allergen.org database: Residue numbering is based on the structure (PDBid:1L6HA) from rice, since there are no solved structures for allergen nsLTP2 proteins. **(a)** The nsLTP1Motif (C3/C50/R44/D43) finds a congruent configuration (C3/C35/R47/D42) in the nsLTP2 structure. While, D42 is conserved in all three sequences, R47 is shifted by one residue. This could be an artifact of the alignment software (both MAFFT [54] and ClustalW [57] have the same alignment). **(b)** Manual insertion of a single gap shows R47 is conserved.

### Structural superimposition based on matching motifs

The structures of nsLTP1 (PDBid:1FK0A) and nsLTP2 (PDBid:1L6HA) have been superimposed using DECAAF [40] (Fig 3). The atoms superimposed were C4/C52/R46 and C35/C3/R47 from nsLTP1 and nsLTP2, respectively (numbering based on corresponding PDBs). The superimposition demonstrates the congruence in R44/D43 in nsLTP1 and R47/D42 in nsLTP2, already highlighted through a pairwise comparison (Table 2). Note, that the global structural configuration of nsLTP1 and nsLTP2 is not homologous with respect to these residues. The cysteines in the disulphide bond is used as a reference point for D43/R43, and is not part of an epitope.

**Figure 3:**
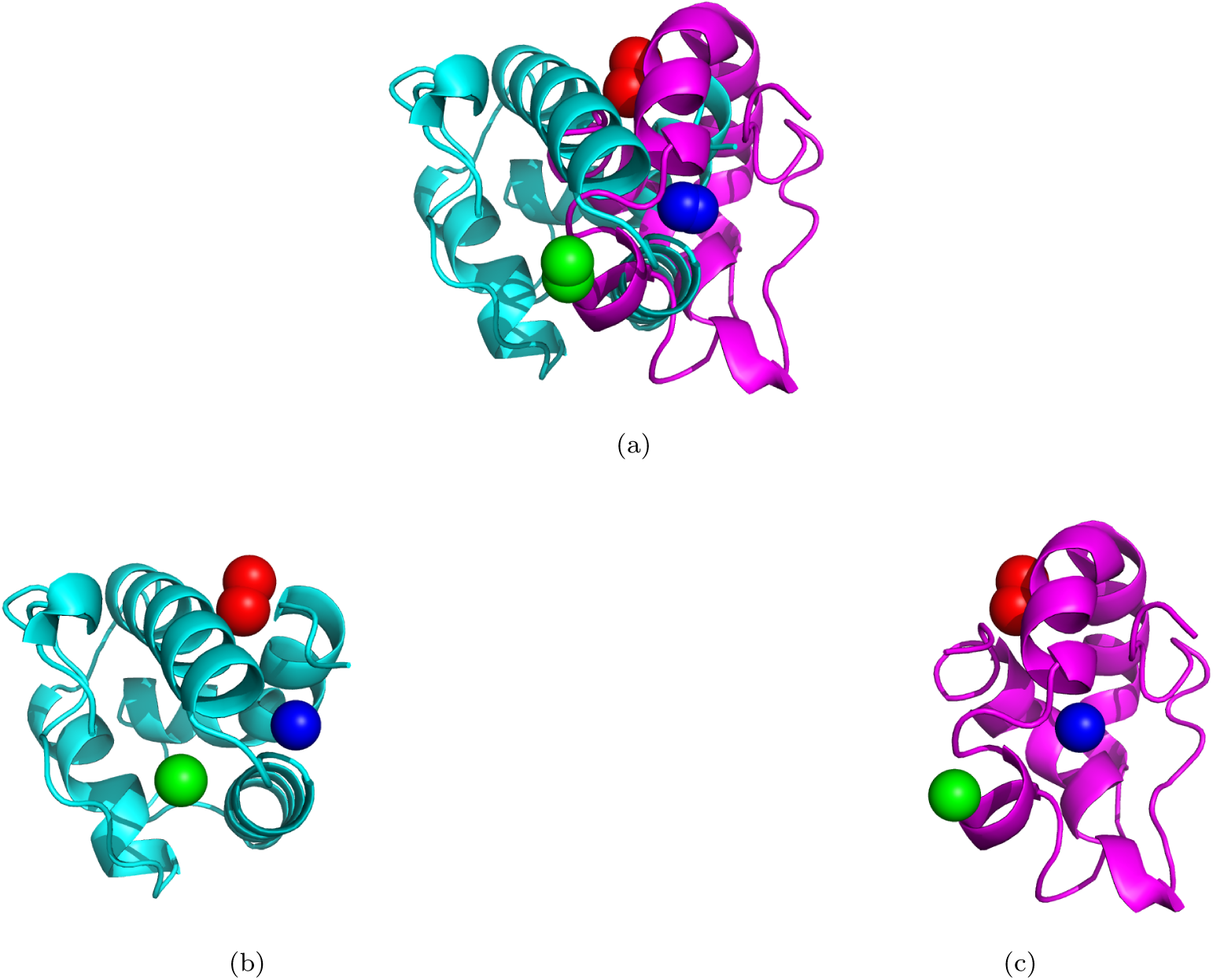
Superimpostion of nsLTP1 (PDBid:1FK0A, in cyan) and nsLTP2 (PDBid:1L6HA, in magenta) using DECAAF: Cysteine residues are in red. **(a)** The atoms superimposed were C4/C5/R46 and C35/C3/R47 from nsLTP1 and nsLTP2, respectively. **(b)** nsLTP1 (PDBid:1FK0A, in cyan). R46 is in green, and D45 is in blue. **(c)** nsLTP2 (PDBid:1L6HA, in magenta). R47 is in green, and D42 is in blue.

### *α*-helices in nsLTP1 and nsLTP2

DSSP analysis identifies secondary structures in proteins [41]. *α*-helices (AH) in nsLTP1 (PDBid:2ALGA) and nsLTP2 (PDBid:1L6HA) are shown in Table 3. The number of AHs in nsLTP1 is five (and not four as mentioned in [22, 42]) (2ALGA.dssp in Dataset1). There is a small 3_10_ AH from residues 11-13, which visually appears to be a single AH (SI Fig 2).

**Table 3:** *α*-helices (AH) in nsLTP1 (PDBid:2ALGA) and nsLTP2 (PDBid:1L6HA): The number of AHs in nsLTP1 is five (and not four as mentioned in [22]). There is a small 3*π* AH from residues 11-13, which visually appears to be a single AH (SI Fig 2). See 2ALGA.dssp in Dataset1 for the DSSP analysis of AH. The basic nature of these nsLTPs is evident from the number of charged residues. HM: Hydrophobic moment, RPNR: Relative proportion of positive residues among charged residues, Len: length of the *α*, NCH: number of charged residues.

| Type | PDB | α-helix | Residues | Len | HM | RPNR | NCH |
| --- | --- | --- | --- | --- | --- | --- | --- |
| nsLTP1 | 2ALGA | 1 | 3 - 10 | 8 | 2.1 | -1 | 0 |
|  |  | 2 | 14 - 19 | 6 | 1.9 | 1 | 1 |
|  |  | 3 | 25 - 37 | 13 | 5.8 | 1 | 1 |
|  |  | 4 | 41 - 57 | 17 | 2 | 0.7 | 3 |
|  |  | 5 | 63 - 72 | 10 | 2.8 | 1 | 1 |
| nsLTP2 | 1L6HA | 1 | 8 - 15 | 8 | 1.1 | -1 | 0 |
|  |  | 2 | 23 - 39 | 17 | 3.3 | 1 | 1 |
|  |  | 3 | 45 - 48 | 4 | 2.5 | 1 | 1 |

### Possible issues with annotations in http://allergen.org/

The site http://allergen.org/ is regularly curated [2]. Possible omissions and mis-annotations are noted here.

1. Keyword search is not uniform - a ‘nsLTP’ search gives 22 and a ‘lipid’ search gives 44 matches. Each keyword misses out on certain entries. There 49 unique matches when these are combined. There are five entries that do not have a tag “lipid” (Table 1 marked with an asterisk).
2. Fragmented allergen (Ole e 7, Uniprot id:P81430, length=21) is not homologous to any other nsLTP, and is probably mis-annotated.
3. PDBid:4XUW is not annotated for the corresponding hazelnut allergen Cor a 8.

### Bioinformatic evaluation of allergenicity in transgenic food

The potential allergenicity of ‘intractable’ proteins [43] expressed in transgenic or GM crops are assessed *in silico* by two criteria: a) > 35% identity over 80-amino-acid stretches and (b) 8-amino-acid contiguous matches [44]. The 8aa window was found to be insu cient in predicting protein allergenicity, in the absence of further homology [45] Here, a more specific search for allergen nsLTPs has been suggested, given the fact that there are many groups of nsLTPs that are non allergens. Furthermore, since there are a limited set of allergens proteins [3], the current work could be easily extended to have a more specific bioinformatic search for allergenicity.

### Conclusions and future work

nsLTPs are an important class of allergens [38]. Although nsLTPs are classified into several groups based on the cysteine residue spacing [19–21], only two classes of these are allergens (nsLTP1 and nsLTP2) [22]. Furthermore, the number of nsLTP1 classified as allergens is much higher than those classified as nsLTP2 (although this can be an artifact of nsLTP2 not been sampled enough times). These two classes have several di erentiating features - sequences, molecular weights, pattern of the conserved disulphide bonds and volume of the hydrophobic cavity [19]. Here, unifying features of nsLTP1 and nsLTP2 responsible for allergenicity is proposed, based on sequence, structural and electrostatic properties. The conservation of R44 in all full length nsLTP1 (only Par j 2 from sticky-weed has an equivalent K44) is in agreement with previous results that this residue is an epitope [23, 24]. Furthermore, D43 is also significantly conserved in most full-length nsLTP1s - only two allergens (Par j 1 and 2) lack that residue, and have possible compensating glutamic residues in the sequence vicinity, missing in sequences that have D43. The presence of a congruent sca old in the nsLTP2 with known structure (PDBid:1L6HA, R47 and D42), and sequence conservation of R47 and D42 in nsLTP2 further strengthens the hypothesis that these are the epitopes responsible for IgE binding. Future work will involve

1. Solving the structure of a nsLTP2 allergen and verification of R47 as an epitope through IgE binding.
2. Solving the structure of the two allergens Par j 1/2 from *P. judaica* which lack D43.
3. Genome wide identification of nsLTPs from other plants commonly causing allergy (walnut [46], chickpea [47], saffron [48]) with these allergen motifs, followed by *in vitro* characterization of these nsLTPs.
4. Determining whether other classes of nsLTP, apart from nsLTP1 and nsLTP2, are allergens.
5. Find whether cross reactivity correlates with structural and/or electrostatic congruence.
6. A similar analysis done in the current work for other classes of allergens.

## Materials and methods

The CLASP algorithm has been detailed previously [29]. In summary, a signature encapsulating the spatial and electrostatic properties are extracted from a given a set of residues, which could be an epitope or catalytic residues, from a protein with known structure. This signature is used to search for congruent matches in a query protein, generating a score which reflects the likelihood that the allergenicity or catalytic activity in the reference protein exists in the query protein. APBS (v1.4) [50, 51] parameters were set as described previously in [29]. APBS writes out the electrostatic potential in dimensionless units of kT/e where k is Boltzmann’s constant, T is the temperature in K and e is the charge of an electron. All protein structures were rendered by PyMOL(TM) Molecular Graphics System, Version 1.7.0.0. (http://www.pymol.org/). *α*-helices and *β*-sheets were extracted using DSSP 2.2.1 [41]. The hydrophobic moment [52] has been computed using PAGAL 1.0 [53]. Protein structures have been superimposed using DECAAF 1.0 [40]. Multiple sequence alignment was done using MAFFT (v7.123b) [54], and figures generated using the ENDscript 2.0 server [55]. In order to obtain a multiple sequence alignment with a single representative of a stereochemical group (positive, negative, aromatic) the following substitutions were done: E>D, K>R, T>S, W>F, Y>F. A grouping algorithm (YeATS-GROUP) was added to the YeATS suite [34]. For a given set of sequences, a BLAST database is created [56]. Each sequence is BLAST’ed to this database, and is linked to another sequence if the BLAST bitscore (BBS) value is more than the specified cutoff (60 in this case). Finally, a group is created such that any sequence in the group has at least one sequence with which it has a homology >BBS=60. The BLAST bitscore was used as a comparison metric instead of the Evalue since it allows di erentiation for high homologies where Evalue goes to zero. Hardware requirements are very modest - all results here are from a simple workstation (8GB ram) and runtimes were a few minutes at the most.

